# The *cnf1* gene is associated to an expanding *Escherichia coli* ST131 *H30*Rx/C2 sublineage and confers a competitive advantage for host colonization

**DOI:** 10.1101/2021.10.13.464032

**Authors:** Landry Laure Tsoumtsa Meda, Luce Landraud, Serena Petracchini, Stéphane Descorps-Declere, Emeline Perthame, Marie-Anne Nahori, Laura Ramirez Finn, Molly A. Ingersoll, Rafael Patiño-Navarrete, Philippe Glaser, Olivier Dussurget, Erick Denamur, Amel Mettouchi, Emmanuel Lemichez

**Affiliations:** Institut Pasteur, Université de Paris, CNRS UMR2001, Unité des Toxines Bactériennes, 75015 Paris, France; Université de Paris, IAME, UMR1137, INSERM, 75018 Paris, France; Université de Paris, 75006, Paris, France; Institut Pasteur, Université de Paris, Bioinformatics and Biostatistics Hub, 75015 Paris, France; Institut Pasteur, Mucosal Inflammation and Immunity group, 75015 Paris, France; EERA Unit “Ecology and Evolution of Antibiotic Resistance”, Institut Pasteur - Assistance Publique/Hôpitaux de Paris - University Paris-Saclay, UMR 3525 CNRS, Paris, France; Institut Pasteur, Yersinia Research Unit, 75015, Paris, France; AP-HP, Laboratoire de Génétique Moléculaire, Hôpital Bichat, 75018 Paris, France

## Abstract

Epidemiological projections point to acquisition of ever-expanding multidrug resistance (MDR) by *Escherichia coli*, a commensal of the digestive tract acting as a source of urinary tract pathogens. We performed a high-throughput genetic screening of predominantly clinical *E. coli* isolates from wide geographical origins. This revealed a preferential distribution of the Cytotoxic Necrotizing Factor 1 (CNF1)-toxin encoding gene, *cnf1*, in four sequence types encompassing the pandemic *E. coli* MDR lineage ST131. This lineage is responsible for a majority of extraintestinal infections that escape first-line antibiotic treatment and has known enhanced capacities to colonize the gastrointestinal tract (GIT). Statistical modeling uncovered a dominant global expansion of *cnf1-*positive strains within multidrug-resistant ST131 subclade *H*30Rx/C2. Despite the absence of phylogeographical signals, *cnf1*-positive isolates adopted a clonal distribution into clusters on the ST131-*H*30Rx/C2 phylogeny, sharing a similar profile of virulence factors and the same *cnf1* allele. Functional analysis of the *cnf1*-positive clinical strain EC131GY ST131-*H*30Rx/C2, established that a *cnf1*-deleted EC131GY is outcompeted by the wildtype strain in a mouse model of competitive infection of the bladder while both strains behave similarly during monoinfections. This points for positive selection of *cnf1* during UTI rather than urovirulence. Wildtype EC131GY also outcompeted the mutant when concurrently inoculated into the gastrointestinal tract, arguing for selection within the gut. Whatever the site of selection, these findings support that the benefit of *cnf1* enhancing host colonization by ST131-*H*30Rx/C2 in turn drives a worldwide dissemination of the *cnf1* gene together with extended spectrum of antibiotic resistance genes.

## INTRODUCTION

CNF1 is a paradigm of bacterial deamidase toxins activating Rho GTPases ^1–4^. Clinical studies document a higher prevalence of the *cnf1*-encoding gene in uropathogenic strains of *Escherichia coli* (UPEC), which belong to the larger group of extraintestinal pathogenic *E. coli* (ExPEC), as compared to commensals from healthy patients ^5–7^. Urinary tract infections (UTI) are common infections that affect more than 150 million individuals annually and are the second cause of antibiotic prescribing ^8^. Despite clinical evidence of a role for *cnf1* in urovirulence ^6^, attempts to define fitness advantages conferred by this toxin in mouse models of UTI have led to opposing conclusions, although these studies do suggest that CNF1 toxin activity may worsen inflammation and tissue damage ^9–13^. Moreover, in an animal model of bacteremia, CNF1 exerts a paradoxical avirulent effect antagonized by the action of the genetically-associated alpha-hemolysin, further blurring the role of CNF1 in host-pathogen interactions ^14–16^. In *E. coli*, there are three types of CNF-like toxins sharing high amino acid sequence identities ^17–20^. However, isolates expressing the CNF2 and CNF3 toxins are rarely detected in extraintestinal infections in humans. Largescale population genetics studies to analyse the distribution of *cnf*-like toxin genes in *E. coli* would give important insights regarding their dynamics within the *E. coli* population.

*E. coli* represents the predominant aerobic bacteria of the gut microbiota, as well as an extraintestinal opportunistic pathogen ^21,22^. Carriage of ExPEC in the gut is a putative source of extraintestinal infections, including UTIs ^23–26^. Only a few sequence types (STs) within the *E. coli* population account for more than half of all *E. coli* strains responsible for extraintestinal infections not causally related to antibiotic resistance ^21,27^. The globally disseminated *E. coli* ST131 has emerged as the predominant lineage responsible for worldwide dissemination of *bla*_CTX-M-15_ extended spectrum beta-lactamase and the rise of multidrug resistant (MDR) extraintestinal infections ^28,29^. This well-defined clonal group is structured into three different clades, with the fluoroquinolone (FQ)-resistant clade C strains subdivided into two subclades comprised of *H*30R/C1 and the dominant expanding *H*30Rx/C2, frequently carrying *bla*_CTX-M-15_ ^30–32^. Enhanced interindividual transmission and dispersal of *E. coli* ST131 lineage likely accounts for the lack of phylogeographical signal ^33^. A larger sampling of strains from the domestic and wild animal world is necessary to better appreciate host specific marks on the evolutionary history of this lineage.

One reason for the unprecedented success of *E. coli* ST131-*H*30 clade C may be its intrinsic capacity to persist in the gastrointestinal tract (GIT) in competition with other strains of *E. coli* ^24,34–37^. Enhanced colonization capacities of the gastrointestinal tract by *E. coli* ST131 likely promote inter-individual transmission, favoring its dissemination in the human population and other hosts, as compared to other lineages ^24,38,39^. The remarkable fitness of this lineage strongly supports the idea of a step-wise acquisition of factors promoting gut colonization, potentially scattered in the UPEC populations. In this case, virulence can be considered as a by-product of commensalism, “virulence factors” being in fact selected for increasing fitness in the commensal niche ^40^.

To better appreciate *cnf1* dynamics, we performed a large-scale screen of the toxin gene distribution in the *E. coli* population. Its increasing prevalence in the ST131-*H*30Rx/C2 lineage led us to test whether an advantage is conferred by *cnf1* for GIT colonization. Wildtype EC131GY from ST131-*H*30Rx/C2 outcompeted the mutant when concurrently inoculated into the gastrointestinal tract, arguing for selection within the gut. The *cnf1*-deleted EC131GY is also outcompeted by the wildtype strain during competitive infection of the bladder. However, in monoinfections both strains infected similarly, pointing to possible positive selection mechanism for *cnf1* during UTI and demonstrating that *cnf1* is not an urovirulence factor. These findings support that the benefit of *cnf1* enhancing host colonization by ST131-*H*30Rx/C2 in turn drives a worldwide dissemination of this lineage.

## RESULTS

### Analysis of the distribution of *cnf* genes in a large collection of *E. coli* genomes

At the start of this study, we mined large genomic datasets from EnteroBase to gain more insight into the distribution of the *cnf1* gene and its close homologs in the population of *E. coli* ^41^. EnteroBase represents an integrated software environment widely used to define the population structure of several bacterial genera, including pathogens. Quantitative information on the collection of 141,234 *E. coli* genomes deposited in EnteroBase are reported in the supplementary figure 1. This collection, starting from 1900, aggregates genomes from strains collected worldwide, but mainly in Europe and North America, and from a wide range of sources but principally human isolates (Sup. Figure 1A, 1B, 1C). Using a Hidden Markov Model (HMM) approach, coupled to amino acid pairwise distance calculation, we retrieved *cnf*-like positive strains and characterized each type of *cnf* sequence. In total, we identified *n*=6,411 *cnf*-positive strains (4.5% of all *E. coli* isolates) with a remarkable dominance of *cnf1* (87.8%, *n*=5,634), as compared to *cnf2* (8.6%, *n*=554) and *cnf3* (3.5%, *n*=223). These strains displayed only one CNF*-*like toxin encoding gene. The prevalent *cnf1* gene in this genomic dataset was widely distributed among isolates of all origins but most notably in the groups denoted humans (5.4% of *n*=48,518 human isolates) and companion animals (24.1% of *n*=2,652 companion animal isolates) (Sup. Figure 1C).

We next studied the distribution of *cnf1* among *E. coli* phylogenetic groups and sequence types (STs). The *cnf1* gene is preferentially associated with isolates from the phylogroup B2, representing 24.3% of *n*=22,305 retrieved genome sequences (Sup. Figure 1D). We observed a tight association of *cnf1* with the most frequently encountered ExPEC sequence types (STs) (Table 1). Notably, a majority of the 5,634 *cnf1*-positive strains segregated among the four sequence types: ST131 (24.5% of *cnf1*-positive strains, *n*=1,382), ST73 (23.2%, *n*=1,308), ST12 (12.4%, *n*=699) and ST127 (10.7%, *n*=601) with the remaining 29.2% of *cnf1*-positive strains widely distributed among 266 other STs. Interestingly, we noticed a steady increase of the percentage of *cnf1-*positive strains in the *E. coli* ST131 lineage from 13% in 2009 up to 23% in 2019 (Figure 1), while this percentage fluctuated around high values in ST73, ST12 and ST127. This analysis reveals a close association of *cnf1* with common ExPEC lineages and a surprising convergent distribution of *cnf1* in ST131, ST73 and ST127 that are representative of adherent-invasive *E. coli* (AIEC) associated with colonic Crohn’s disease and known to have enhanced capacities to colonize the gastrointestinal tract ^21,42,43^.

**Figure 1:**
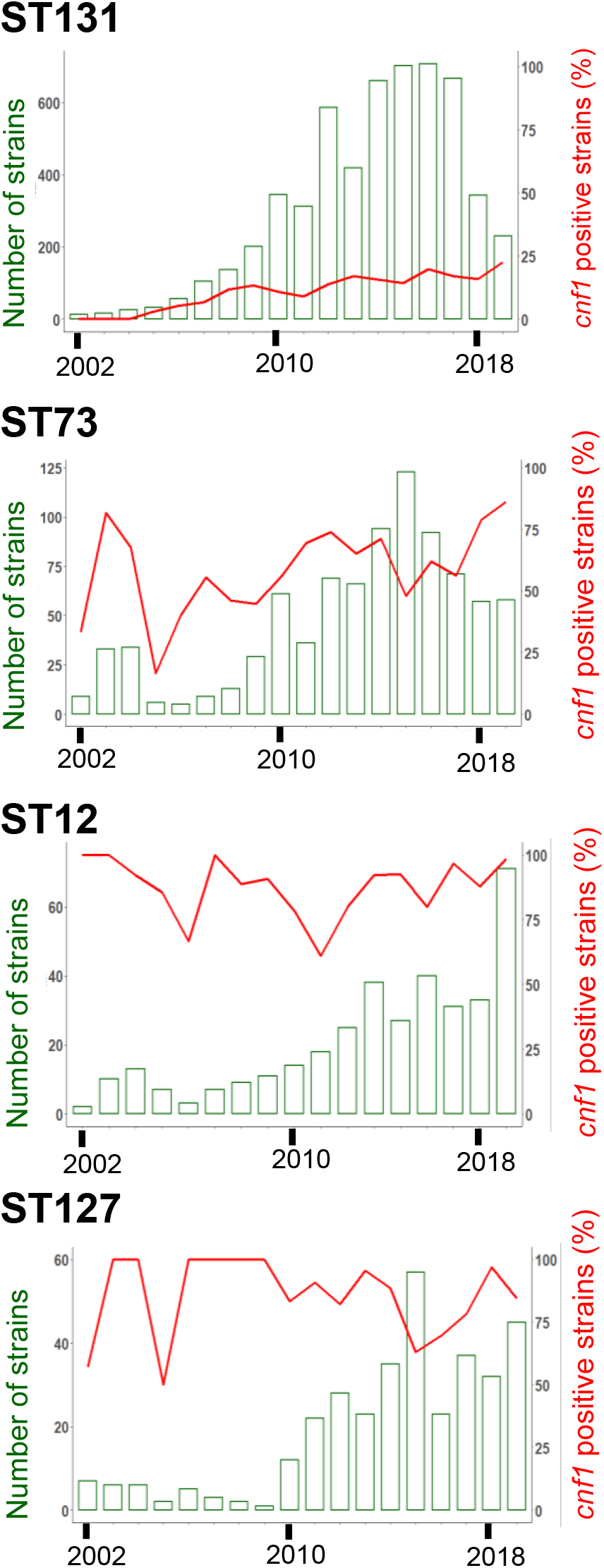
Prevalence overtime in representative *E. coli* sequence types bearing *cnf1*. Bar chart show number of *E. coli* strains from ST131, ST127, ST73 and ST12 isolated each year during the period 2002-2019, left y-axis. Percentages of *cnf1*-positive strains per year, right y-axis.

**Table 1:** Distribution of phylogroups and sequence types among *E. coli cnf*-positive strains from EnteroBase. The total number and the percentage of each phylogroup and most dominant sequence types (STs) among *cnf*-positive strains are indicated

| Phylogroups | ST | Number of strains |  |  |  |  | Percentage of Phylogroup or Sequence type in CNF-positive strains |  |  |
| --- | --- | --- | --- | --- | --- | --- | --- | --- | --- |
|  |  | All | CNF+ | CNF1+ | CNF2+ | CNF3+ | CNF1 | CNF2 | CNF3 |
| A | Total A | 34,982 | 51 | 0 | 28 | 23 | 0 | 5.05 | 10.31 |
|  | ST10 | 8,748 | 24 | 0 | 17 | 7 | 0.0 | 3.1 | 3.1 |
|  | ST342 | 325 | 16 | 0 | 0 | 16 | 0.0 | 0.0 | 7.2 |
| B1 | Total B1 | 37,262 | 527 | 96 | 373 | 58 | 1.7 | 67.3 | 26.0 |
|  | ST101 | 938 | 93 | 24 | 69 | 0 | 0.4 | 12.5 | 0.0 |
|  | ST392 | 79 | 66 | 0 | 66 | 0 | 0.0 | 11.9 | 0.0 |
|  | ST58 | 1,487 | 44 | 9 | 35 | 0 | 0.2 | 6.3 | 0.0 |
|  | ST29 | 496 | 35 | 0 | 0 | 35 | 0.0 | 0.0 | 15.7 |
|  | ST2217 | 46 | 31 | 0 | 31 | 0 | 0.0 | 5.6 | 0.0 |
|  | ST5738 | 24 | 23 | 0 | 23 | 0 | 0.0 | 4.2 | 0.0 |
|  | ST21 | 5,082 | 10 | 0 | 0 | 10 | 0.0 | 0.0 | 4.5 |
|  | ST343 | 134 | 2 | 0 | 0 | 2 | 0.0 | 0.0 | 0.9 |
|  | ST2836 | 63 | 2 | 0 | 0 | 2 | 0.0 | 0.0 | 0.9 |
|  | ST4063 | 3 | 2 | 0 | 0 | 2 | 0.0 | 0.0 | 0.9 |
| B2 | Total B2 | 22,305 | 5,478 | 5,414 | 63 | 1 | 96.1 | 11.4 | 0.4 |
|  | ST131 | 9,242 | 1,383 | 1,382 | 0 | 1 | 24.5 | 0.0 | 0.4 |
|  | ST73 | 2,071 | 1,308 | 1,308 | 0 | 0 | 23.2 | 0.0 | 0.0 |
|  | ST12 | 809 | 699 | 699 | 0 | 0 | 12.4 | 0.0 | 0.0 |
|  | ST127 | 709 | 601 | 601 | 0 | 0 | 10.7 | 0.0 | 0.0 |
|  | ST372 | 366 | 206 | 206 | 0 | 0 | 3.7 | 0.0 | 0.0 |
|  | ST95 | 1,882 | 173 | 147 | 26 | 0 | 2.6 | 4.7 | 0.0 |
|  | ST141 | 360 | 164 | 164 | 0 | 0 | 2.9 | 0.0 | 0.0 |
|  | ST998 | 175 | 149 | 149 | 0 | 0 | 2.6 | 0.0 | 0.0 |
|  | ST80 | 152 | 109 | 105 | 4 | 0 | 1.9 | 0.7 | 0.0 |
|  | ST537 | 50 | 35 | 35 | 0 | 0 | 0.6 | 0.0 | 0.0 |
|  | ST647 | 28 | 26 | 0 | 26 | 0 | 0.0 | 4.7 | 0.0 |
| C | Total C | 3,465 | 56 | 45 | 10 | 1 | 0.8 | 1.8 | 0.4 |
| D | Total D | 9,905 | 37 | 20 | 13 | 4 | 0.4 | 2.3 | 1.8 |
| E | Total E | 16,391 | 155 | 7 | 14 | 134 | 0.1 | 2.5 | 60.1 |
|  | ST11 | 13,639 | 113 | 0 | 0 | 113 | 0.0 | 0.0 | 50.7 |
|  | ST5592 | 5 | 5 | 0 | 0 | 5 | 0.0 | 0.0 | 2.2 |
|  | ST11457 | 4 | 4 | 0 | 0 | 4 | 0.0 | 0.0 | 1.8 |
| F | Total F | 2,957 | 38 | 37 | 0 | 1 | 0.7 | 0.0 | 0.4 |
| G | Total G | 1,862 | 34 | 0 | 34 | 0 | 0.0 | 6.1 | 0.0 |
|  | ST117 | 1,383 | 31 | 0 | 31 | 0 | 0.0 | 5.6 | 0.0 |
| Clade I | Total CI | 406 | 18 | 0 | 18 | 0 | 0.0 | 3.2 | 0.0 |
|  | ST3057 | 41 | 11 | 0 | 11 | 0 | 0.0 | 2.0 | 0.0 |
| Clade II | Total CII | 6 | 0 | 0 | 0 | 0 | 0.0 | 0.0 | 0.0 |
| Clade III | Total CIII | 39 | 0 | 0 | 0 | 0 | 0.0 | 0.0 | 0.0 |
| Clade IV | Total CIV | 39 | 0 | 0 | 0 | 0 | 0.0 | 0.0 | 0.0 |
| Clade V | Total CV | 166 | 0 | 0 | 0 | 0 | 0.0 | 0.0 | 0.0 |
|  | Other 358 STs | 34,599 | 1,044 | 803 | 215 | 26 | 14.3 | 38.8 | 11.7 |

### *cnf1*-positive strains segregate into monophyletic groups in ST131 phylogeny

The rising prevalence of *cnf1* in *E. coli* ST131 motivated us to study its distribution in this lineage, as its phylogenetic structure is well defined and displays a major FQ-resistant clade largely independent of geographical signal ^30–33^. EnteroBase contained 9,242 genomes of *E. coli* ST131 at the time of analysis (November 2020). To ease genomic analysis, we retained 5,231 genomes that were isolated from 1967 to 2018. We built a Maximum Likelihood phylogenetic tree based on a total of 37,304 non-recombinant SNPs. Phylogenetic distribution of strains showed an expected dominant population of clade C (76%, *n* = 3,981; 99% *fimH*30), as compared to clade A (11%, *n* = 569; 92% *fimH*41) and B (13%, *n* = 68; 62% *fimH*22) (Figure 2A, detailed in Sup. Figure 2A). We also found an expected co-distribution of *parC* (S80I/E84V) and *gyrA* (S83L/D87N) alleles that confer strong resistance to FQ in most strains from clade C (99.84%, *n*=3,975 strains), and a tight association of the *bla*_CTX-M-15_ ESBL gene (85%, *n*=2,194 isolates) with strains from subclade *H*30Rx/C2. The high number of strains gave enough resolution to distinguish two sublineages, C2_1 and C2_2, originating from C2_0 (Figure 2A). From available metadata, we verified the absence of overall geographical and temporal links in the phylogenetic distribution of *E. coli* ST131 strains (Sup. Figure 2B). In conclusion, large scale phylogenetic reconstruction of ST131 genomes from EnteroBase showed an expected phylogenetic distribution within clades and subclades of genetic traits defining this lineage.

**Figure 2:**
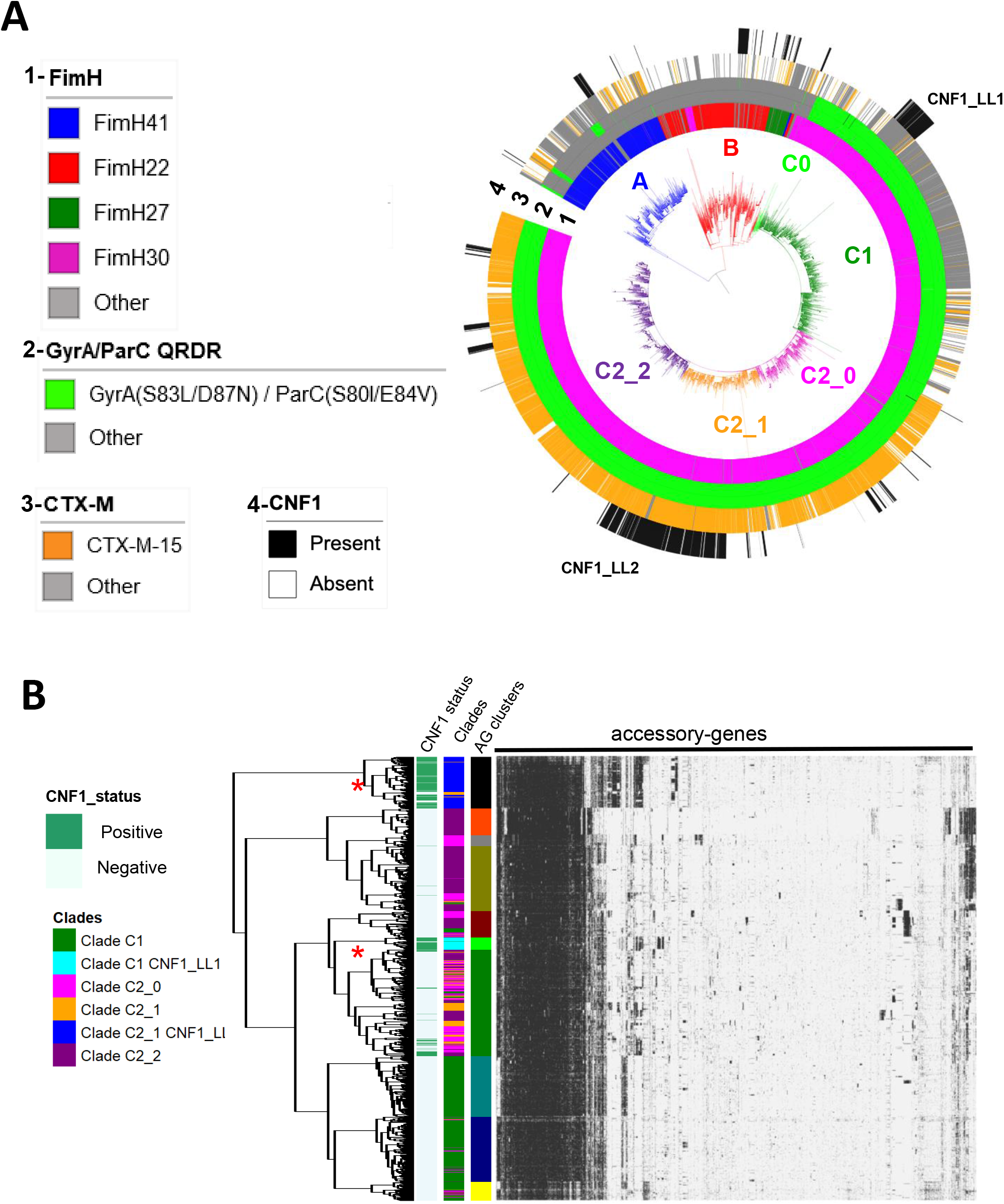
Dynamic of CNF1-encoding gene in *E. coli* ST131 from EnteroBase. **A)** Maximum likelihood phylogeny of *E. coli* ST131 from EnteroBase (Sup. Figure 2 for extended information). The phylogeny was constructed with 5,231 genomes for a total of 37,304 non-recombinant core-genome SNPs. The different clades and subclades A, B, C0, C1, C2_0, C2_1, C2_2 are highlighted in blue, red, light green, green, pink, orange and purple respectively. From inside to outside circles are indicated (1) *fimH* alleles, (2) *gyrA* and *parC* alleles conferring resistance to FQ (shown in green), (3) strains positive for *bla*_CTX-M-15_ (shown in orange) and (4) strains bearing *cnf*1 gene (shown in black). **B)** Hierarchical clustering of strains from clade C (*n* = 3981 strains) based on their accessory gene content. The pan-genome is composed of 51,742 genes including 2,672 genes that are present in 98% of the strains. The graph displays the 7,678 genes identified as present in at least 50 and less than 3,930 genomes. The colored annotation indicates (from left to right) the presence of *cnf1* (CNF1_status), clades (C1, C1 CNF1_LL1, C2_0, C2_1, C2_1 CNF1_LL2, C2_2) and accessory genes cluster (AG_clusters). Large lineages of *cnf1*-positive strains in clades C1 and C2_1 are denoted CNF1_LL1 and CNF1_LL2, respectively.

We next analyzed the distribution of *cnf1*-positive strains (*n*=725) in *E. coli* ST131 phylogeny (Figure 2A, black stripes). The *cnf1*-positive strains were preferentially associated with clade C2 (*n*=520), as compared to clade C1 (*n*=101), clade B (*n*=72) and clade A (*n*=32) (Figure 2A). Strikingly, most *cnf1*-positive strains segregated into lineages in all clades and subclades with a noticeable distribution of *cnf1*-positive ST131 strains in two large lineages (LL) in *H*30R/C1 (*n*=101 *cnf1-*positive strains/107 strains in CNF1_LL1) and in *H*30Rx/C2_1 (*n*=396 *cnf1-* positive strains/425 strains in the CNF1_LL2) (Figure 2A). We then analyzed the diversity of alleles of *cnf1* to define their distribution in ST131 phylogeny (Sup. Table 1). A similar analysis was performed with the alpha-hemolysin encoding gene, *hlyA*. We found a wide co-distribution of one combination of alleles of *cnf1* (allele P1_*cnf1*_, 85,1%) and alpha-hemolysin encoding gene *hlyA* (allele P1_*hlyA*_, 77,2%) in *E. coli* ST131 clade A and C, whereas strains from clade B displayed a large range of combinations of various alleles (Sup. Figure 2A). Together, our data point to a clonal expansion of worldwide disseminated ST131-*H*30 strains having the same allele of *cnf1*. Together, this prompted us to perform a clustering analysis of ST131-*H*30 strains according to their accessory gene contents. We generated a pan-genome matrix of 51,742 coding sequences from the *n*=3,981 strains of clade C. The dataset of accessory genes was built from *n*=7,678 sequences that were present in at least 50 and no more than 3,931 strains. We conducted a hierarchical clustering of strains according to the Ward’s minimum variance-derived method ^44^ and retained 10 distinct accessory gene clusters. Strikingly, this revealed a conservation between phylogenetically-defined groups CNF1_LL1 and CNF1_LL2 and groups defined by their accessory gene contents (Figure 2B). Indeed, the hierarchical clustering was most evident for CNF1_LL2, showing a differential enrichment of *n*=1,434 genes as compared to other strains from clade C, determined with Scoary (Bonferroni-adjusted *P*-value <0.05) ^45^. Together, these data point towards intensive group-specific diversification of accessory gene content in *cnf1*-positive clusters in ST131-*H*30.

### *cnf1*-positive strains of *E. coli* ST131 segregate between two clade-specific virulence profiles

We then defined strain contents in virulence factors (VF) and acquired antibiotic-resistance genes (RG) to perform an unbiased analysis of their distribution into clusters, using a latent block model approach. Acquired antibiotic-resistance genes in ST131 genomes were identified with ResFinder ^46^. Profiles of virulence factors were defined with the database published by Petty and colleagues ^31^. The unsupervised clustering procedure retained a total of 10 RG-clusters and 7 VF-clusters (Figure 3A). Differences in number of VFs and RGs between clusters were all significant (Figure 3B). We found that *cnf1*-positive strains were scattered among several RG clusters (Figure 3A, left panel). By contrast, most *cnf1*-positive strains segregated into the cluster VF4 (84% of *cnf1*-positive strains, *n*=609) with the remaining 16% strains being distributed between VF1 (15%) and other VF clusters (1%) (Figure 3A, right panel). In contrast to RG-clusters, we observed that VF-clusters formed phylogenetically defined groups (Figure 3C). A majority of *cnf1*-positive strains from clade A and B were positive for the VF1 cluster, whereas *cnf1*-positive strains from clade C were positive for the VF4 cluster. With a mean value of 33 virulence factors (Figure 3B), VF4-positive strains displayed the largest arsenal of virulence factors. The VF1 profile was more specifically defined by the presence of genes encoding the IbeA invasin and IroN Salmochelin siderophore receptor (Sup. Figure 3A). By contrast, major determinants of the VF4 cluster encompassed *cnf1* and *hlyA* (54% and 61% in VF4 versus 34% in VF1 and 3% in all other VFs). Specific VF determinants of VF4 also encompassed genes encoding the UclD adhesin that tipped F17-like chaperone-usher (CU) fimbriae cluster and PapG II adhesin from pyelonephritis-associated pili (pap) operon (Sup. Figure 3A) ^47,48^. These elements can be genetically associated and constitute the backbone of *cnf1*-bearing pathogenicity islands (PAI) II_J96_ from the O4:K6 *E. coli* strain J96, although PAI II_J96_ contains a *papG* class III sequence (Sup. Figure 3B). In good agreement, analysis of several complete sequences of *cnf1*-bearing PAI II_J96_-like from ST131-*H*30 showed a conservation of a module containing this set of genes, defining VF4 (Sup. Figure 3B).

**Figure 3:**
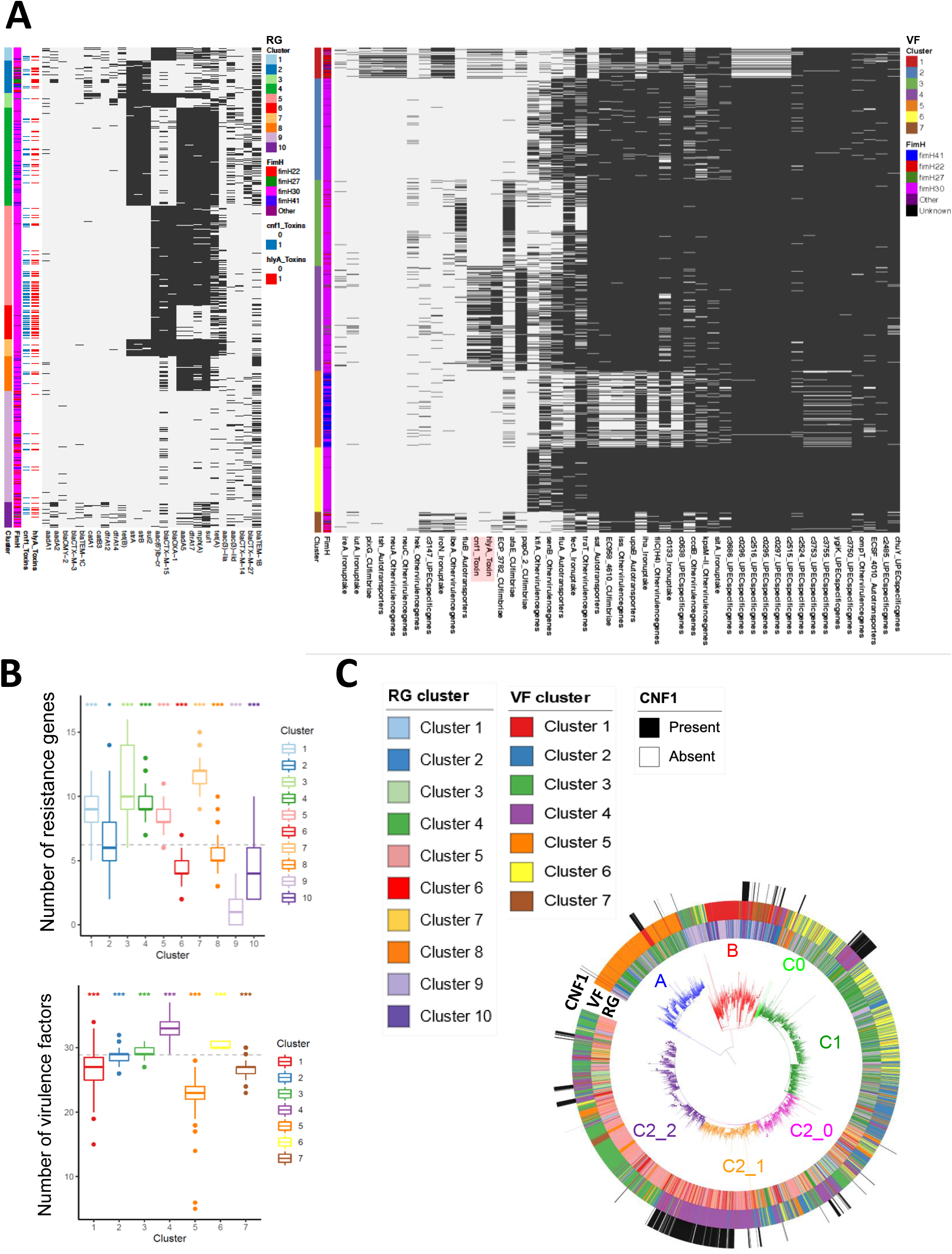
Co-clustering of acquired antibiotic-resistance gene and virulence factors in *E. coli* ST131. **A)** Heatmaps show clusters of antibiotic acquired-resistance gene (RG) (left panel) or virulence gene (VF) (right panel) profiles (Sup. table 2) constructed using a binary latent block model between strains by row and RGs or VFs by column. Black lines indicate the presence of RG or VF in each strain. Annotations are displayed on the right of each heatmap: information about strain clusters and *fimH* alleles together with *hlyA* and *cnf1* carriage. **B)** Box-and-whisker plot showing the distribution of strains according to their content of acquired antibiotic-resistance genes (upper panel) or content of virulence factors (lower panel). The dotted line shows the mean number of RG or VF. All one-versus-all comparisons of VF and RG contents between clusters (\**P* < 0.05, \*\*\**P* < 0.001). **C)** RG, VF clusters and *cnf1* carriage are displayed on the *E. coli* ST131 phylogenetic tree. The different clades and subclades A, B, C0, C1, C2_0, C2_1, C2_2 are highlighted in blue, red, light green, green, pink, orange and purple respectively.

### *cnf1*-positive strains display dominant expansion in ST131-*H*30Rx/C2

We next analyzed the temporal distribution of *cnf1-*positive strains within clades and subclades. Using a Generalized Linear Models (GLM) approach, we first verified within our dataset the increase of *fimH*30-positive isolates over time (clade C) in *E. coli* ST131 that was maximal in *H*30Rx/C2 (*P*<2 10^−16^) (Figure 4A). We also noted a significant increase in the proportion of *cnf1*-positive strains over time in *E. coli* ST131 (Figure 4B, top panel). The GLM was then fitted on years, clades, and subclades. We tested the significance of the year effect and *P*-values were corrected for multiple comparisons using Tukey’s method. The year effect was not significant for clade A, B, or subclade *H*30R/C1 (Figure 4B). Instead, we observed a significant increase of the proportion of *cnf1*-positive strains within *H*30Rx/C2 over time (*P*=1.25 10^−11^). In addition, the GLM fitted curves predicted that the prevalence of *cnf1*-positive strains within *H*30Rx/C2 sublineage would be approximately 50% (confidence interval of 95% [43% to 58%] in 2018; [47% to 64%] in 2019). Predictive values were confronted to the prevalence of *cnf1* in ST131 strains isolated in 2018 or 2019 in a second independent dataset up-loaded from EnteroBase in September 2020. This confirmed the rising prevalence of *cnf1*-positive strains within the sublineage *H*30Rx/C2 up to 45% in 2018 and 48% in 2019. In conclusion, we identified a dominant expansion of *cnf1*-positive strains within ST131-*H*30Rx/C2.

**Figure 4:**
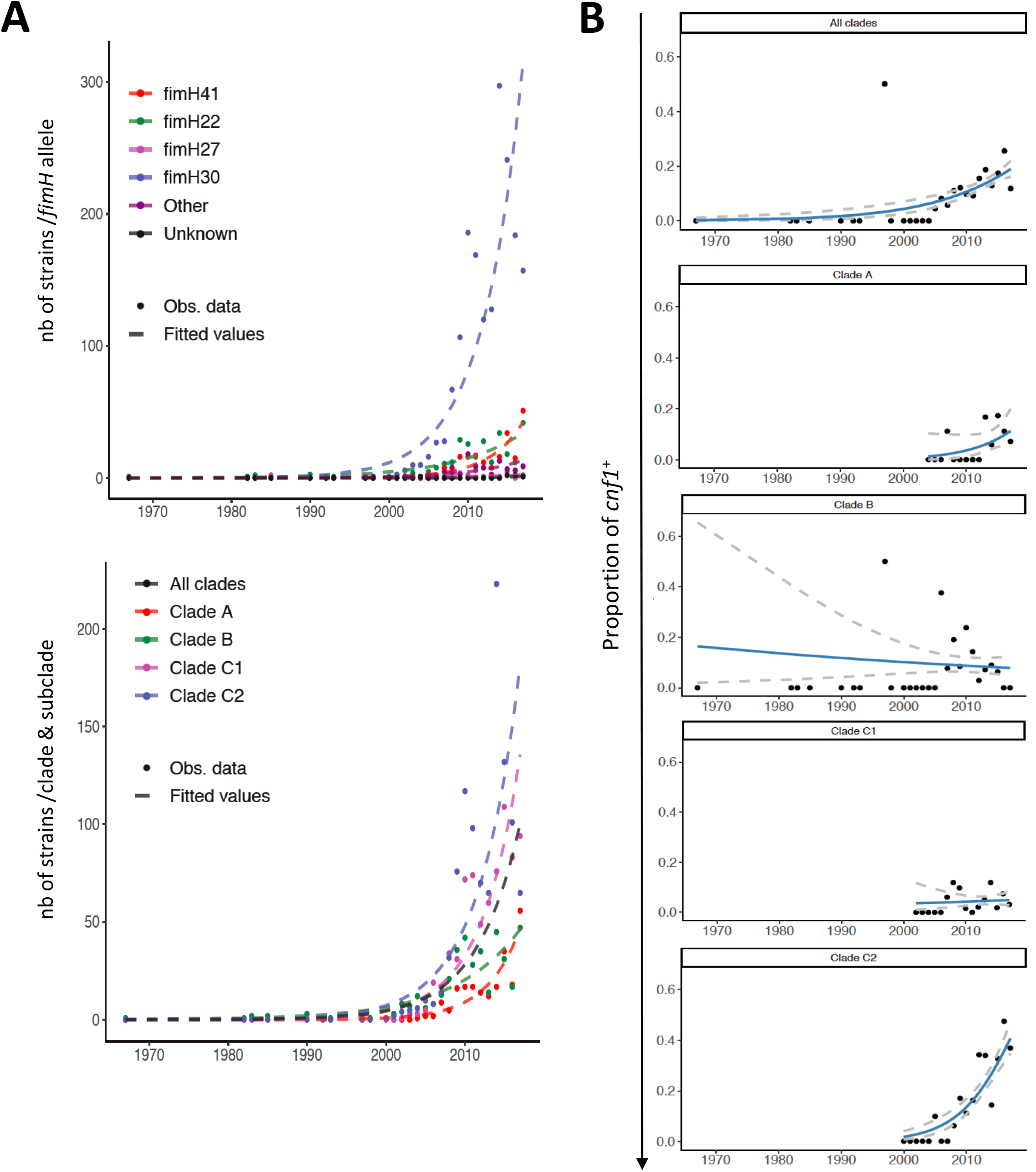
Increase over the year in the proportion of *cnf1*-positive strains in *E. coli* ST131 H30Rx/C2. **A)** Distribution of *fimH* alleles (upper panel) or clades/subclades (lower panel) within the study population of *E. coli* ST131. Both figures show observed counts per year (dots) and data fitted lines (dashed lines) with a generalized linear model (Poisson regression). **B)** Increase of the proportion of *cnf1*-positive strains in the whole *E. coli* ST131 population along time (top panel, *P* = 7.41 10-7) and by clades and subclades. The black dots represent the observed proportion of *cnf1*-positive strains by year with fitted line of a logistic regression model (blue curves). Dashed grey lines display the 95% confidence intervals. The *P*-values are not significant for clade A (*P* = 0.287), B (*P* = 0.952), *H*30R/C1 (*P* = 0.992) and significant for *H*30Rx/C2 (*P* = 1.25 10^−11^).

### *cnf1* confers a competitive advantage for bladder infection and gut colonization in a ST131-*H*30Rx/C2 strain

The dominant expansion of *cnf1*-positive strains in ST131 *H*30Rx/C2 prompted us to explore whether CNF1 confers a competitive advantage for bladder infection and/or intestinal colonization. In the cohort SEPTICOLI of bloodstream infections in human adults ^49^, we identified a VF4/*cnf1*-positive strain of *E. coli* ST131 *H*30Rx/C2, here referred to as EC131GY (Sup. Figure 4). This strain is amenable to genetic engineering and displays a *cnf1*-bearing PAI (PAI II_EC131GY_) highly similar to the prototypic PAI II_J96_ from the J96 (O4:H5:K6) UPEC strain (Sup. Figure 3B) ^50^. We generated a EC131GY strain in which *cnf1* was replaced with a kanamycin resistance cassette (EC131GYΔ*cnf1::kan*^r^) and verified the absence of CNF1 expression (Sup. Figure 5A). We next verified, *in vitro*, the absence of fitness cost due to the kanamycin resistance cassette as shown by equal growth of parental and Δ*cnf1::kan*^r^ EC131GY strains, and the absence of competition between the strains when grown together (Sup. Figure 5B and 5C). Considering the tight association of *cnf1* with clinical strains of *E. coli* responsible for UTI, we first investigated the impact of the toxin during concurrent infection of the bladder with wild-type EC131GY and EC131GYΔ*cnf1::kan*^r^. Wild-type *E. coli* outcompeted the isogenic *cnf1*-deficient EC131GY in the first 24 hours, when bacteria must rapidly establish their niche in the face of passive and innate immune host defenses (Figure 5A). This fitness advantage was maintained at day 3 and 7, demonstrating that *cnf1* plays a role in the early stages of UPEC pathogenesis, as previously suggested ^9^. No difference of colonization of wild-type EC131GY and EC131GYΔ*cnf1::kan*^r^ was observed in monomicrobial bladder infections (Figure 5B). This finding can be interpreted as a positive selection mechanism to maintain the CNF1 gene during UTI, considering that a loss of *cnf1* would be detrimental for bacterial fitness in a mixed population. We then explored the impact of *cnf1* in GIT colonization, again by competitive infection with EC131GY WT and EC131GYΔ*cnf1::kan*^r^, using intra-gastric gavage ^51^. Longitudinal measurements of CFU in the feces showed that CNF1 conferred an advantage to wild-type EC131GY over the EC131GYΔ*cnf1::kan*^r^ isogenic strain for gut colonization from 9 days after oral gavage, which persisted over 27 days (Figure 5B). Together, these data uncover the advantage conferred by CNF1 in a setting of competitive UTI and for intestinal colonization by the VF4/*cnf1*-positive EC131GY strain from the ST131-*H*30Rx/C2 lineage.

**Figure 5:**
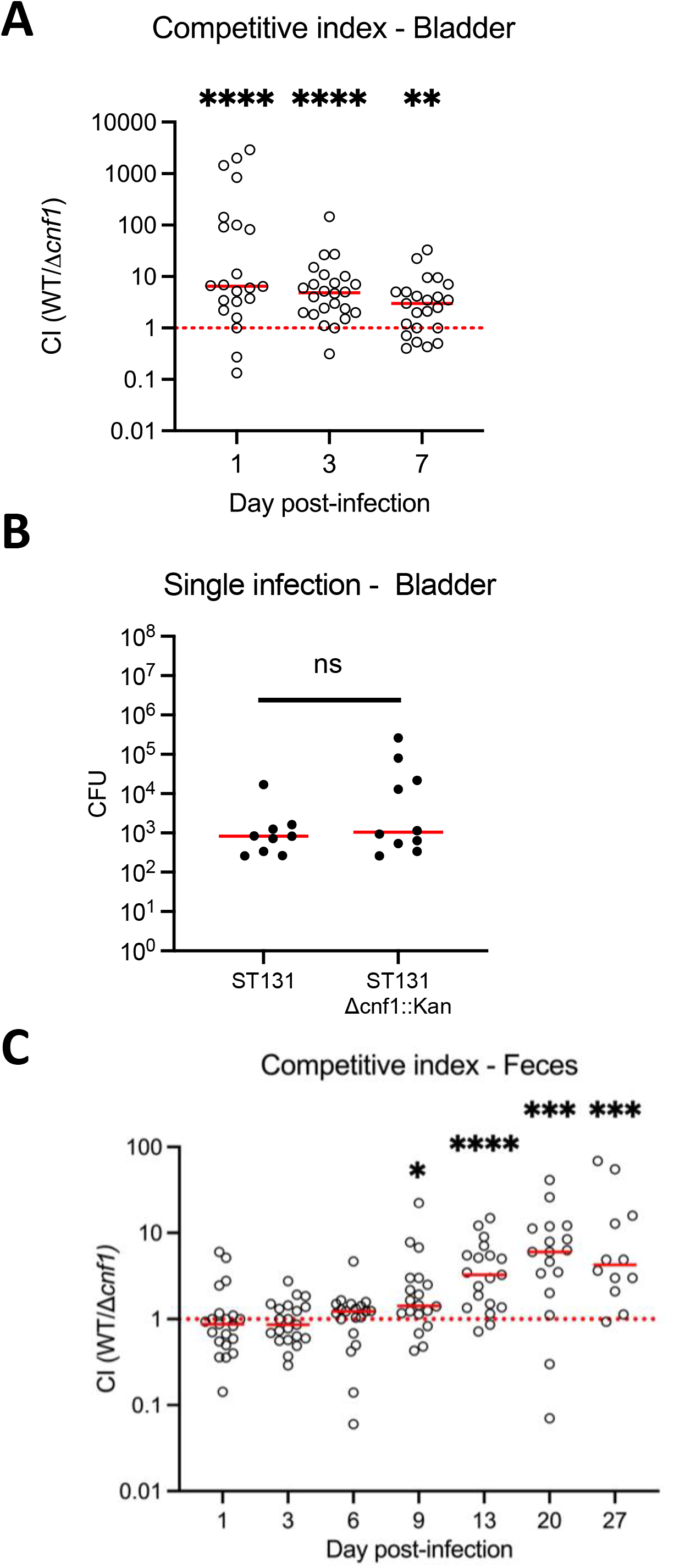
CNF1 promotes ST131-H30Rx/C2 bladder and intestinal colonization. Mice were infected concurrently **(A)** or separately **(B)** with wild-type EC131GY (WT) and EC131GY Δ*cnf1*::kan^r^ (Δ*cnf1*) via intravesical instillation of the bladder. For GIT colonization, mice were pretreated with streptomycin and subsequently infected concurrently via the oral route with EC131GY WT and Δ*cnf1* **(C)**. Levels of viable bacteria in bladder homogenates or feces were assessed at indicated times by measuring colony forming units (CFU). Data represent the competitive index (CI) (A and C) or CFU per bladder (B) for each animal and medians (red bar). Total of *n*=15-18 (bladder CI, three replicates), *n*=9-10 (bladder single, two replicates at day 1) and *n*=21 (intestine, three replicates). **\****P* < 0.05, **\*\****P* < 0.01, **\*\*\****P* < 0.001, **\*\*\*\****P* < 0.0001 and ns : non-significant by Wilcoxon signed-rank test.

## DISCUSSION

Initially thought to be absent in the *Escherichia coli* ST131 lineage, the *cnf1* gene was estimated to be found in approximately 15% of this lineage, among 99 isolates from distinct geographical locations across the world, in 2014 ^31,52^. Large-scale genetic analysis of more than five thousand isolates of *E. coli* ST131 from EnteroBase, a database widely used by clinicians, gives here sufficient statistical power to unveil a dominant expansion trend of *cnf1*-positive strains within clade *H*30Rx/C2. Our analysis supports the hypothesis of a recent expansion of a large phylogenetic subcluster of *cnf1*-positive ST131-*H*30Rx/C2 strains circulating between humans and dogs ^53,54^. In addition, we document a stable population dynamic of *cnf1*-positive *H*30R/C1 strains within clade C1. This raises the question of whether *cnf1* confers a fitness advantage at the population level. Our compelling findings ascribed such a feature of *cnf1* to specific genetic backgrounds, thereby enhancing the expansion and dissemination of a subpopulation of ST131-*H*30Rx/C2 within the ST131 lineage. Furthermore, we report the high prevalence of *cnf1* gene in the three sequence types ST73, ST12 and ST127 of *E. coli* that have different antibiotic resistance profiles. Together, this points to a role of *cnf1* in the dynamics of ExPEC that is independent from antibiotic resistance genetic backgrounds. The rising prevalence of *cnf1*-positive *H*30Rx/C2, and evidence of their mobilization between humans and dogs ^53^, suggest that *cnf1* enhances the dissemination of *H*30RxC2 within households with companion animals, which is likely driven by an increased ability to compete for GIT colonization. In further support of this conclusion, we found a prevalence of 24% of *cnf1*-positive strains in the group companion animals from the EnteroBase database. Finally, we report a high occurrence of the *cnf1* gene in common AIEC pathotypes responsible for Crohn’s disease and known to colonize the GIT well ^21,42,43^. These findings highlight the importance of studying the interplay between CNF1 and the gut mucosa for persistence and inflammatory bowel diseases.

The competitive advantage conferred by *cnf1* during the acute phase of UTI (i.e., 24 hours) suggests this toxin promotes FimH-dependent invasion of urothelial cells, which results in the formation of intracellular bacterial communities (IBCs) ^8,55^. In support of this hypothesis, cell biology studies show that CNF1 promotes invasion of host cells by *E. coli* through its capacity to activate host Rho GTPases ^20,56–58^. Although this remains to be formally demonstrated, CNF1 deamidase likely exacerbates the activation of Rho GTPases, which are required for type I pili-mediated host cell invasion ^59^. Importantly, in contrast to concurrent infection, *cnf1* confers no detectable virulence advantage during bladder monoinfection. Considering that UTI caused by *E. coli* are usually dominated by one strain, we propose that the fitness advantage conferred by *cnf1* during concurrent infection could reflect a positive selection mechanism to maintain the gene during UTI. Alternatively, as *cnf1* also confers a fitness advantage in the gut commensal niche which is the primary *E. coli* habitat, the selective pressure occurs in the gut and *cnf1* confers virulence as a by-product of commensalism ^40^. This mechanism has been shown for the PAIs of the B2 ST127 strain 536 ^60^. The F17-like pilus adhesin UclD from *cnf1*-bearing PAI confers a competition advantage for gut colonization, while it shows no virulence role in UTI ^51^. Therefore, this also points for *cnf1-*driven positive selection as a potential broader mechanism to maintain the PAI during UTI.

Our findings that *cnf1* gives a competitive advantage for GIT colonization also raise the interest of defining epistatic relationships between factors encoded within the core set of genes of the PAI II_EC131GY_ from ST131 H30Rx/C2 for colonization and bacterial persistence in tissues. Indeed, these operons encode F17-like pili, the P-fimbriae tipped with PapG class II adhesin, and the *hlyA* toxin, as well as a gene encoding haemagglutinin *in E. coli* K1 (Hek) ^61–63^. This also includes elements of oxidative stress adaptation, namely the methionine sulfoxide reductase complex MsrPQ encoding genes *yedYZ*, which may work against CNF1-generated oxidative stress ^64,65^.

Collectively, our findings point towards a bidirectional interplay between *cnf1* and the *E. coli* ST131 lineage to enhance host colonization by *H*30Rx/C2 whatever the site of selection and to promote a worldwide dissemination of the Cytotoxic Necrotizing Factor 1-encoding gene together with extended spectrum of antibiotic resistant genes.

## MATERIAL and METHODS

### *E. coli* genome collection

Collection of 141,234 *E. coli* genome sequences from EnteroBase (November 2020) (http://enterobase.warwick.ac.uk) ^41^. Strain’s metadata (collection year, continent, source niche of isolation and sequence type) were also retrieved (Sup. Table 3). Assemblies were downloaded in GenBank format and proteomes generated using annotations provided in GenBank files.

### *In silico* detection and typing of CNF-like toxin encoding genes

The search for *cnf* genes in *E. coli* genomes was carried out with a domain specific Hidden Markov Models (HMM) profile built with 16 representative sequences of CNF1 catalytic domain (Sup. Table 4) using HMMER (http://hmmer.org/) ^66^. Protein sequences from positive hits were extracted from EnteroBase annotated *E. coli* proteomes and submitted to Clustal Omega for the computation of pairwise distances of the sequences, along with representative sequences of CNF-like toxin (CNF1 (AAA85196.1), CNF2 (WP_012775889.1) and CNF3 (WP_02231387.1)). Distances were used to determine the type of toxin with a threshold value of 0.1. In total 2.7% of HMM-positive sequences with a threshold value above 0.1 against all type of CNF-like toxin or below 0.1 against at least two type of CNF-like toxin were excluded from the analysis.

### ST131 dataset structure and phylogenomic analysis

The database used for phylogenetic and statistical analyses consists of whole-genome sequences of *E. coli* ST131 isolates collected by mining EnteroBase from 1967 to 2018 ^41^. Leaning on Find ST(s) tool from EnteroBase, we retained a total of 5,231 genome assemblies and associated metadata, including information of the isolation date, country and source of isolates (Sup. table 5). Phylogeny of ST131 isolates was resolved using core non-recombinant SNPs defined with Parsnp (in total 37,304 SNPs) ^67^ and Gubbins v2.3.4 ^68^. A maximum-likelihood tree was then estimated with RAxML v8.2.8 applying a general time-reversible substitution-model with a gamma distribution rate across sites and with an ascertainment bias correction ^69^ and the resulting tree was edited with the interactive Tree of Life (iTol) v4 program ^70^.

### *In silico* antimicrobial resistance and virulence-associated markers

GyrA and ParC protein sequences were retrieved from the EnteroBase annotated genomes, and aligned with the mafft L-INS-I approach ^71^. After a visual inspection of the alignment, in-house customized perl scripts (https://github.com/rpatinonavarrete/QRDR) were used to identify the amino acids at the quinolone resistance-determining region (QRDR) (positions 83 and 87, and 80 and 84 in GyrA and ParC, respectively). Search for *cnf1* and *hlyA* alleles in ST131 genomes dataset was carried out by Blastn analysis. Sequences were next aligned with Muscle ^72^ and curated to remove incomplete sequences. SNPs were then extracted using SNP-sites ^73^. To determine strain specific VF profiles, annotated VFs from UPEC described in ^31^ were translated and pBLASTed against ST131 genomes dataset considering only hits with e-value < 10^−5^ and identical matches > 95% (sup. Table 2) ^74^. Acquired antibiotic-resistance genes (RGs) in ST131 genomes were defined with ResFinder ^46^.

### Generalized linear model

Proportion of *cnf1* along time was modeled using a generalized linear model (logistic regression) adjusted on the effect of years and clades with an interaction between these two factors. First, to test if the evolution of *cnf1* proportion was either specific to each clade or global, the significance of the interaction term was tested with a likelihood ratio test, which compares the above-mentioned model against the null model, with no interaction. Then, we investigated the possible increase of the proportion of *cnf1* within each clade. The significance of the slope coefficient for each clade was tested by computing contrasts of the above model. *P*-values were adjusted for multiplicity using single-step correction method. The distribution of *fimH* alleles and clades/subclades within the study population of *E. coli* ST131 was analyzed with a similar approach, except that a Poisson regression model was used to model counting data. The hypothesis testing strategy to investigate the significance of the increase of *fimH* alleles and clades/subclades along time is discussed above.

### Co-clustering method

Statistical analyses were performed using R software version 3.6.0. A total of 20 strains from the collection of 5,231 strains of *E. coli* ST131 were removed from the analysis due to incomplete associated metadata. The clustering of strains with specific virulence or acquired antibiotic-resistance gene profiles was performed with binary latent block model, implemented in the R package blockcluster ^75^. In this package, the model, a mixture of Bernoulli distributions proposed by ^76^, is estimated using an efficient EM algorithm. As proposed by the authors, the number of clusters was estimated by maximizing the ICL criterion on a bidimensional grid of parameters making this unsupervised classification procedure automatic.

### Pan-genome analysis

The pangenome of *E. coli* ST131 was estimated using Roary, a high-speed pan genome pipeline analysis tool ^77^. Roary returns as output, the gene presence/absence matrix. The matrix was curated to retain genes present in at least 50 genomes and less than 3980 genomes (7678 sequences), that constituted our accessory genes pool dataset. Hierarchical clustering analysis was then conducted by using the pheatmap package in R (cran.r-project.org/web/packages/pheatmap/index.html). The gene presence/absence file generated by Roary was further analyzed using Scoary ^45^ with a significant Bonferroni-adjusted P-value < 0.05 for genes associated to *cnf1*-positive lineages (Sup. Table 8).

### Mouse colonization model

Local Animal Studies Committee and National Research Council approved all procedures used for the mouse experiments described in the present study (APAFIS#26133-202006221228936 v1, 2016–0010. For gut colonization, groups of female C57BL/6 mice aged 6–7 weeks (Charles River) were pretreated with a single dose of streptomycin (1 g/kg in 200 μl water) *per os* 1 day prior to gavage, as described in ^51^. The strains derived from the clinical strain H1-001-0141-G-Y, here referred to as EC131GY (de Lastours et al., 2020), are described in the extended materials and methods section. Mice were co-infected *per os* with 2×10^9^ CFU of each strain in 200 μl PBS. Fecal pellets were collected from every individual mouse at indicated times, weighed and homogenized in 500 μl phosphate-buffed saline (PBS) pH 7.2 by vigorous vortexing. CFUs were determined by plating serial dilutions on selective LB agar plates. Strains were prepared for infection as follows: a single colony of EC131GY or its derivative was inoculated in 10 ml selective LB medium and incubated at 37°C under static conditions for 24h. Bacteria were then inoculated in 25 ml fresh selective LB medium at 1:1000 dilution and incubated at 37°C under static conditions for 18-24h. Bacteria were then washed twice in cold PBS, and concentrated in PBS at approximately 2×10^9^ CFU per 200 μl. Inocula titers are verified in parallel for each infection. For intravesical infection: Urinary tract infection was induced in mice as previously described ^78,79^. Briefly, a single colony of EC131GY or the *cnf1* mutant was inoculated in 10 ml LB medium with antibiotics and incubated at 37°C under static conditions for 18h. Mice were infected with a total of 10^7^ CFU of bacteria in 50 μl PBS via a rigid urinary catheter under anesthesia. To calculate CFU, bladders were aseptically removed and homogenized in 1 ml of PBS. Serial dilutions were plated on LB agar plates with antibiotics, as required. The competitive index (CI) was calculated as: CFU WT output strain/CFU mutant output strain, with the verification in each experiment that CFU WT input strain/CFU mutant input strain was close to 1. A Wilcoxon signed-rank test was performed to assess the statistical significance of differences in CI over time. Statistical analyses were performed using GraphPad Prism 9.

## ACKNOWLEDGMENTS

This work was supported by the “Fondation ARC” PJA 20191209650, the “Fondation pour la Recherche Médicale” (Equipe FRM 2016, DEQ20161136698), Ligue Nationale contre le Cancer Subvention de Recherche Scientifique, RS20/75-63 and the French National Research Agency (ANR-10-LABX-62-IBEID, INCEPTION) and ANR-17-CE17-0014. The plasmid pKOBEG was kindly provided by Jean-Marc Ghigo.

## AUTHOR CONTRIBUTIONS

Bioinformatics analyses were performed L.T.M., S.D.-D., R.P.N. and analyzed by E.L., L.L., P.G. and E.D. Statistical analyses were performed by L.L. and E.P. *In vivo* experiments were coordinated by A.M., M.A.I., O.D. and performed by M.-A. N., A.M. and L.R.F. with strains engineered by S.P. and A.M. The research was coordinated by E.L. and manuscript drafted with help of L.T.M., L.L., O.D., E.D. and P.G. Manuscript was reviewed and approved by all authors.

