## Supplementary figures legends for "The *cnf1* gene is associated to an expanding *Escherichia coli* ST131 *H30*Rx/C2 sublineage and confers a competitive advantage for host colonization"

### SUPPLEMENTAL FIGURE LEGENDS

**Sup. Figure 1: Available metadata associated with genomes of *E. coli* from EnteroBase**

**A-B)** Available metadata of *E. coli* isolates from EnteroBase show their distribution according to the period of isolation (A) and geographic origin (B). Red line shows the percentage of E. coli strains that were *cnf*-toxins positive over the years. **C-D)** Tables show the percentage of *cnf1*-positive strains for each origin (C) or each phylogroup (D). Total corresponds to the number of strains with available metadata information.

**Sup. Figure 2: Maximum likelihood phylogenetic tree of *E. coli* ST131 genomes**

**A-B)** Maximum likelihood phylogeny of *E. coli* ST131 from EnteroBase with different clades and subclades A, B, C0, C1, C2_0, C2_1, C2_2 highlighted in blue, red, light green, green, pink, orange and purple, respectively. Phylogeny was constructed with 5,231 genomes for a total of 37,304 non-recombinant core-genome SNPs and visualized with iTol. Colors strips surrounding the phylogram represent, from inside to outside circles: somatic (O) and flagellar (H) antigen combination (1) and alleles of *fimH* (2), *gyrA* (3), *parC* (4), *bla*$_{CTX\text{-}M}$ (5), *hlyA* (6) and *cnf1* (7) for each strain. **B)** geographical origin (1) and period of isolation of each strain (2) of *E. coli* ST131 isolates together with *cnf1*-positive strains from large clusters (3) CNF1_LL1 (green) and CNF1_LL2 strains (blue).

**Sup. Figure 3: *cnf1* together with elements of PAI IIJ96 specify VF4 cluster**

**A)** Left graph shows prevalence of virulence factors from VF4 in *E. coli* ST131 study population (x-axis) and the 1,128 VF4-positive strains (y-axis). Similar analysis for VF1-positive strains (right graph). **B)** Comparison of the genomic organization with PAI II$_{EC131GY}$ from EC131GY and 8 other *E. coli* ST131 strains from EnteroBase (ESC_VA2376AA_AS, ESC_JA0942AA_AS, ESC_VA2412AA_AS, ESC_VA2988AA_AS, ESC_LA1508AA_AS, ESC_VA2411AA_AS, ESC_JA0943AA_AS and ESC_JA0947AA_AS), as well as reference PAI II$_{J96}$ from the strain J96 (GCA_000295775.2). Genomic islands containing *cnf1*-encoding gene were defined with IslandPath-DIMOB of Islandviewer 4. Annotated sequences were aligned and visualized using Easyfig. Red lines between PAIs show > 60% blast homologies. Genes located inside PAIs are displayed as colored arrows together with annotation of large operons. Genes or operon encoding the haemagglutinin from *E. coli* K1 (Hek), fimbrial adhesin PapG, F17-like pili, Cytotoxic Necrotizing Factor-1 (CNF1), alpha-hemolysin (HlyA), yedYZ-encoding methionine sulfoxide reductase (MsrPQ), histidine-kinase two-component system (YedVW), contact-dependent growth inhibition (CdiI), cryptic phage-bearing toxin/antitoxin systems CP4-57 and CP4-44, and lysine decarboxylase (CadCBA) are annotated. Other open reading frames are indicated in blue.

**Sup. Figure 4: Phylogenetic tree revealing the position of EC131GY**

Localization of EC131GY onto the phylogenic tree of C2_2 *E. coli* ST131 strains. Phylogeny was constructed using core-genome non-recombinant SNPs of C2_2 E. coli ST131 strains including EC131GY genome. EC131GY is indicated (red arrow). Numbers indicate information of *fimH allele* (1), *bla*$_{\text{CTX-M-15/14}}$ (2), VF clusters (3), geographic origin (4), source niche (5) and *presence/absence of cnf1* (6) for each strain. Lower inset (*) shows a close-up on EC131GY and surrounding strains.

**Sup. Figure 5: Characterization of EC131GY WT and mutant strain**

**A)** Representative Immunoblots anti-CNF1 and anti-HlyA showing levels of expression of both toxins in EC131GY WT and Δ*cnf1::Kan*$^r$ (Δ*cnf1)*. Immunoblots anti-RNA-Pol show loading control. **B)** Kinetics of individual growth of *E. coli* ST131 EC131GY WT and Δ*cnf1*::Kan$^r$ (EC131GY Δ*cnf1*) monitored at $OD_{600}$. Data show one representative experiment performed with five biological replicates ± SD. **C)** Kinetic of growth competition between *E. coli* ST131 EC131GY WT and Δ*cnf1*::Kan$^r$ mixed 1:1. Bacteria were grown together for 5 hours at 37°C with shaking and CFU/ml determined on LB and LB kanamycin plates from multiple serial dilutions. Each dot corresponds to the competitive index value (CI) between WT and Δ*cnf1*::Kan$^r$ (EC131GYΔ*cnf1*) at indicated time points. Data are shown as mean ± SEM, *n*=3 independent experiments. Data not significant difference by Mann–Whitney U test.

**Sup. Table 1: *cnf1* and *hlyA* SNPs profile in ST131 genomes**

First two files correspond to the list of SNPs in *cnf1* (file1) and *hlyA* (file2) genes of ST131 genomes and distribution in profiles. The red square indicates the presence of a SNP in each profile. File 3 corresponds to the co-distribution of *cnf1* and *hlyA* SNPs profiles in ST131 genomes.

**Sup. Table 2: Profiles of acquired antibiotic-resistance genes and virulence factor encoding genes**

File 1 corresponds to the list of acquired antibiotic-resistance genes (RGs) and virulence factors (VFs) studied and their occurrence in the population of *E. coli* ST131 deposited in EnteroBase. Note that we retained RGs and VFs that show a differential occurrence in genomes, i.e. in less than 5,131 and more than 100 genomes. File 2 corresponds to the distribution of RGs and VFs profile (express as percentage) in strains within RG clusters and VF clusters.

**Sup. Table 3: Metadata describing *E. coli* genomes obtained from EnteroBase**

**Sup. Table 4: List of representative sequences of CNF1 catalytic domain**

**Sup. Table 5: *E. coli* ST131 strains from the dataset and associated metadata**

**Sup. Table 6: List of strains and plasmids used in the study**

**Sup. Table 7: List of primers**

**Sup. Table 8: Scoary results**

### EXTENDED MATERIAL AND METHODS

**Bacterial strains and growth conditions.** The ST131 strain EC131GY (H1-001-0141-G-Y) has been originally isolated from a patient suffering from bacteremia [1]. The strain is naturally resistant to ampicillin. A streptomycin-resistant isolate was selected and used to engineer the *cnf1* mutant strain. EC131GY is susceptible to gentamicin (cmi 0.5 mg/L) and resistant to cefotaxime (cmi >256 mg/L). The *cnf1* mutant was engineered from the streptomycin-resistant strain EC131GY by substitution of the *cnf1* gene by a kanamycin-resistance cassette as detailed below. Bacteria were grown on agar plate at 30°C or 37°C starting from frozen samples kept in -80°C in 30% glycerol. From the fresh agar plate, bacteria were grown overnight in liquid culture under 120 rpm shaking condition in LB broth, at 30°C or 37°C. When needed, ampicillin (100 µg/ml), kanamycin(50 µg /ml), or chloramphenicol (25 µg /ml) were added. Strains and plasmids used in this study are listed in supplementary table 6.

**Construction of gene deletion mutant strains.** Deletion of *cnf1* gene from the chromosome of EC131GY has been performed with the Lambda Red recombination system for gene replacement as described in [2]. Briefly, primers for amplification of the kanamycin cassette and the flanking FRT regions in pKD4 have been designed to target the first and the last 81 nucleotides of the *cnf1* gene (Sup. table 7). The resulting PCR product was purified using commercial kits (Macherey Nagel). The strain carrying the temperature-sensitive helper plasmid pKOBEG coding for the Lamda red recombinase system was treated as in [2]. The resulting mutants were tested for the gene replacement by PCR with primers listed in table 2, and pKOBEG plasmid loss was verified on LB agar plates with chloramphenicol.

**Growth curves.** Growth curves were performed using a FLUOstar Omega microplate reader. Briefly, starting from a fresh overnight culture, bacteria were diluted 1/100 in 5 mL LB supplemented with streptomycin 200µg/ml. 200 µl of each culture were placed as 5 replicates in a 96 flat bottom plate (Greiner) and incubated for 12 hours at 37°C with 120 rpm orbital shaking. Absorbance at 600nm was measured every 10 min.

**Western blot.** Bacterial pellets were collected in RIPA buffer. The lysates were boiled in 1x Laemmli buffer 5 minutes at 100°C and resolved on 8% SDS-PAGE, transferred to nitrocellulose membrane (GE Healthcare). The proteins were colored with ponceau S (Biorad) and the membrane blocked with 5% milk in TBS-T (Euromedex). Membranes were incubated with the primary antibody: CNF1 (Santa Cruz sc52655 clone NG8 1/1000), RNA Polymerase (Biolegend 699907 clone NT73 1/1000) and rabbit serum (1/1000) against the conserved amino acids 914–936 of HlyA, described in [3]. Membranes were washed with TBS-T and incubated with horseradish peroxidase (HRP)-conjugated secondary antibodies for 1h. Signals were observed using Immobion Western Chemiluminescent HRP Substrate (Merck).

**Supplementary table 6: list of strains and plasmids**

| Strains or plasmids | Relevant characteristics | Source | Reference |
|---|---|---|---|
| ***E. coli*** | | | |
| EC131GY | Clinical isolate H1-001-0141-G-Y-StrepR | | 1 |
| ***E. coli* mutant strain** | | | |
| EC131GY*Δcnf1::Kan* | $Kan^R$; *cnf1* deletion via $Kan^R$ insertion | This work | |
| **Plasmids** | | | |
| pKD4 | $Amp^R$; template for PCR of $Kan^R$ | CGSC | 2 |
| pKOBEG | $Cm^R$; Lambda Red recombinase system | Gift from JM. Ghigo | 4 |

CGSC: Coli Genetic Stock Center

**Supplementary table 7: list of primers**

| Primers | |
|---|---|
| **cnf.H1-P1** | 5’AGGTCTCTGTCTGAGAGTTATTCTCTGAATGCAGATGCCTCCGAAATATCGGTATTGAAGGTATTTTCAAAAAAATTTTGA |
| **cnf.P2-H2** | 5’TCAAAATTTTTTTGAAAATACCTTCAATACCGATATTTCGGAGGCATCTGCATTCAGAGAATAACTCTCAGACAGAGACCTGGTCCATATGAATATCCTCCTTAG |
| **cnfver.fw** | 5’ATGGGTAACCAATGGCAACAAAAATATCTT |
| **cnfver.rev** | 5’ATGGGTAACCAATGGCAACAAAAATATCTT |
