## Supplementary figures for "The *cnf1* gene is associated to an expanding *Escherichia coli* ST131 *H30*Rx/C2 sublineage and confers a competitive advantage for host colonization"

A

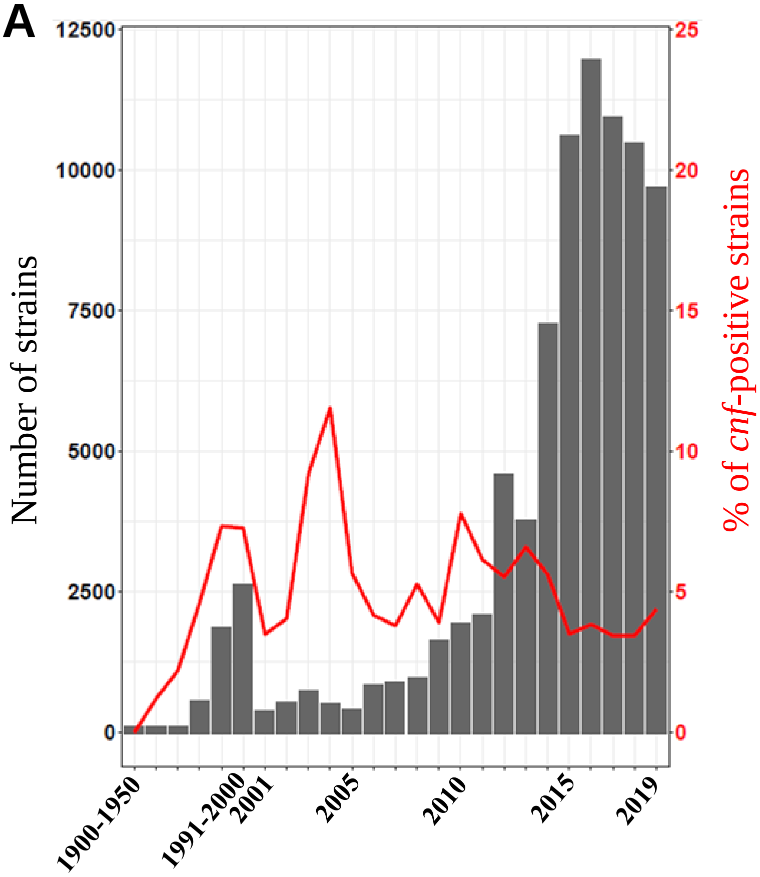


B

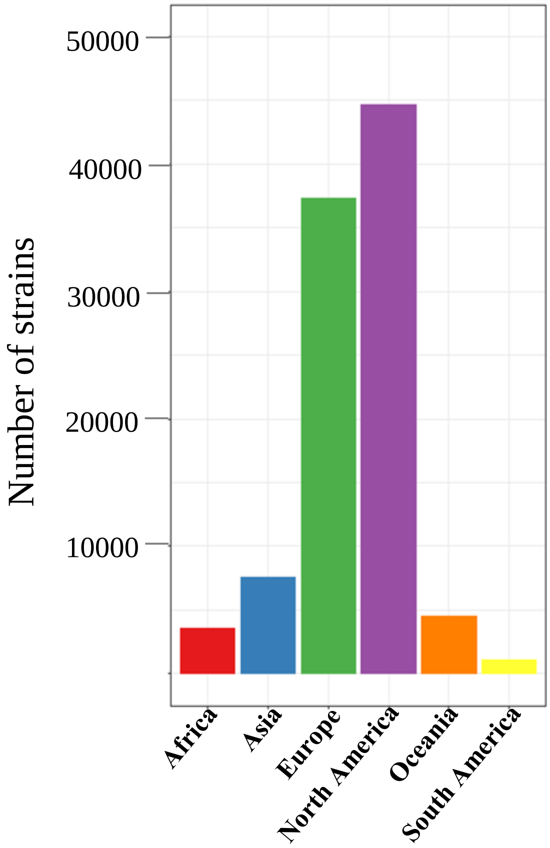


C

| SOURCE NICHE | NEG | POS | TOTAL | *cnf1*+ (%) |
|---|---|---|---|---|
| **Human** | 45875 | 2643 | 48518 | 5,45 |
| **Companion Animal** | 1979 | 629 | 2608 | 24,12 |
| **Animal Feed** | 541 | 7 | 548 | 1,28 |
| **Aquatic Animal** | 400 | 9 | 409 | 2,20 |
| **Environment** | 6300 | 198 | 6498 | 3,05 |
| **Food** | 2227 | 58 | 2285 | 2,54 |
| **Livestock** | 13794 | 319 | 14113 | 2,26 |
| **Poultry** | 7919 | 59 | 7978 | 0,74 |
| **Wild Animal** | 2511 | 141 | 2652 | 5,32 |
| **ND** | 53099 | 2526 | 55625 | 4,54 |

D

| PHYLOGROUPS | NEG | POS | TOTAL | *cnf1*+ (%) |
|---|---|---|---|---|
| **A** | 34982 | 0 | 34982 | 0,00 |
| **B1** | 37166 | 96 | 37262 | 0,26 |
| **B2** | 16891 | 5414 | 22305 | 24,27 |
| **C** | 3420 | 45 | 3465 | 1,30 |
| **D** | 9885 | 20 | 9905 | 0,20 |
| **E** | 16384 | 7 | 16391 | 0,04 |
| **F** | 2920 | 37 | 2957 | 1,25 |
| **G** | 1862 | 0 | 1862 | 0,00 |
| **Clade I** | 406 | 0 | 406 | 0,00 |
| **Clade II** | 6 | 0 | 6 | 0,00 |
| **Clade III** | 39 | 0 | 39 | 0,00 |
| **Clade IV** | 39 | 0 | 39 | 0,00 |
| **Clade V** | 166 | 0 | 166 | 0,00 |

Sup. Fig. 1 : Tsoumtsa et al.

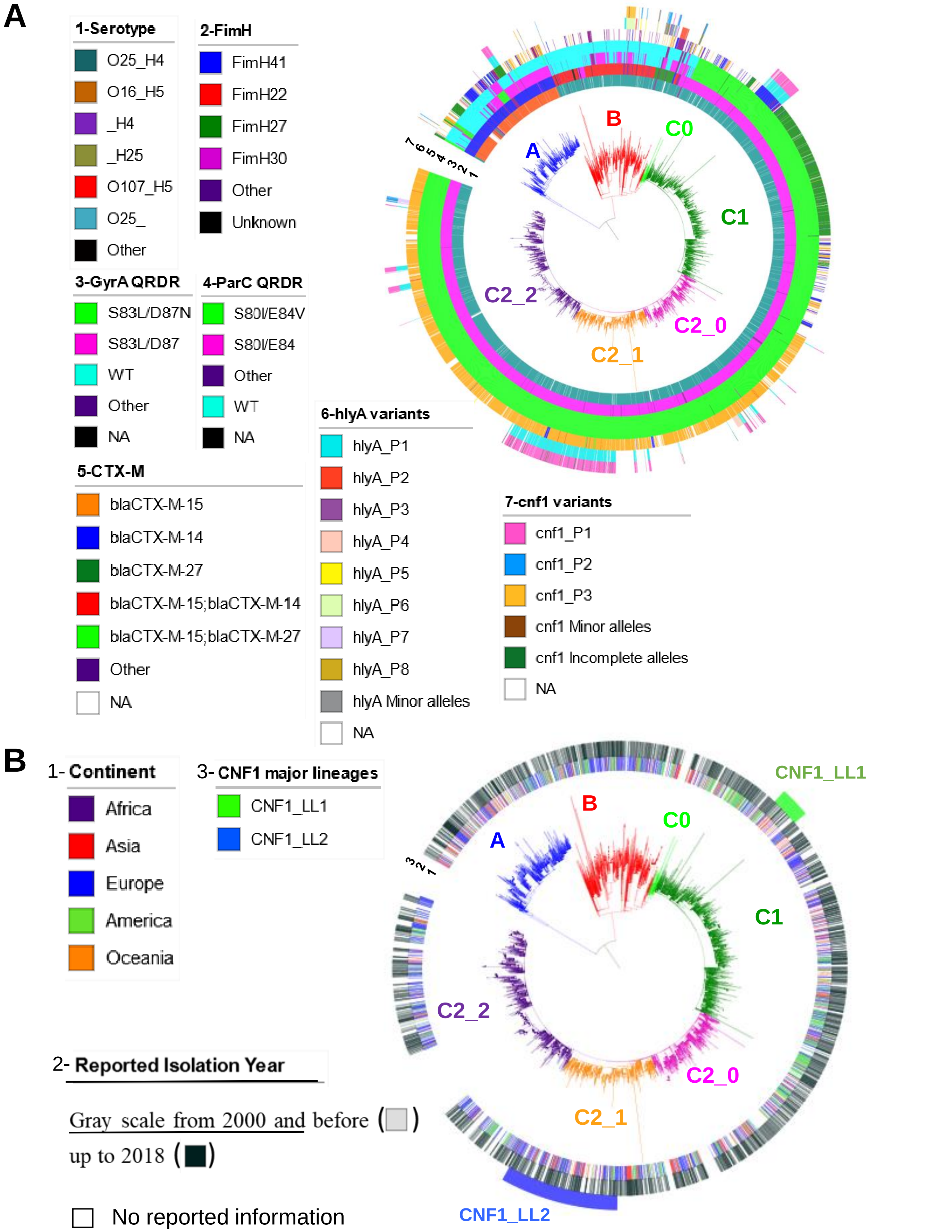


Sup. Fig. 2 : Tsoumtsa et al.

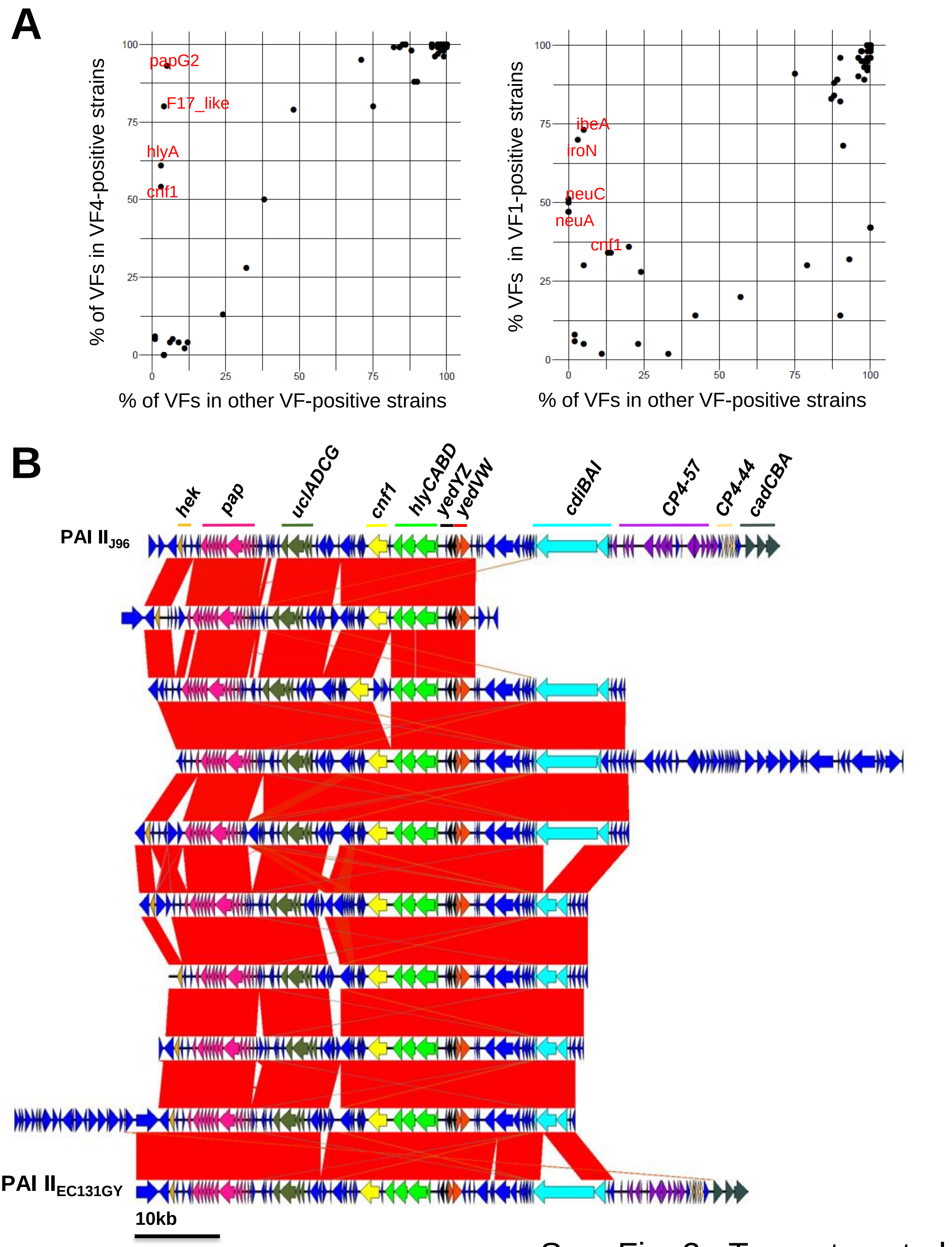


Sup. Fig. 3 : Tsoumtsa et al.

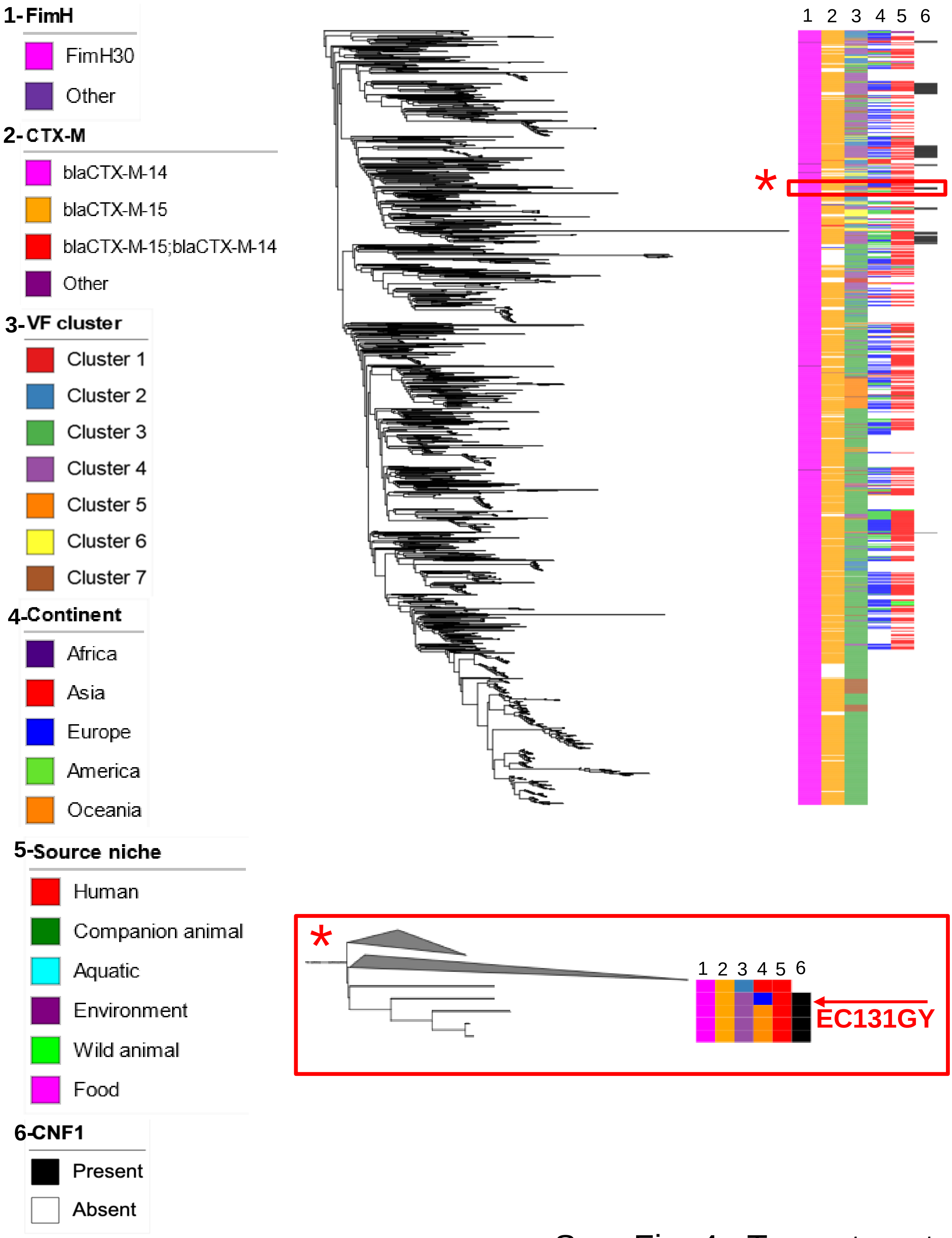


Sup. Fig. 4 : Tsoumtsa et al.

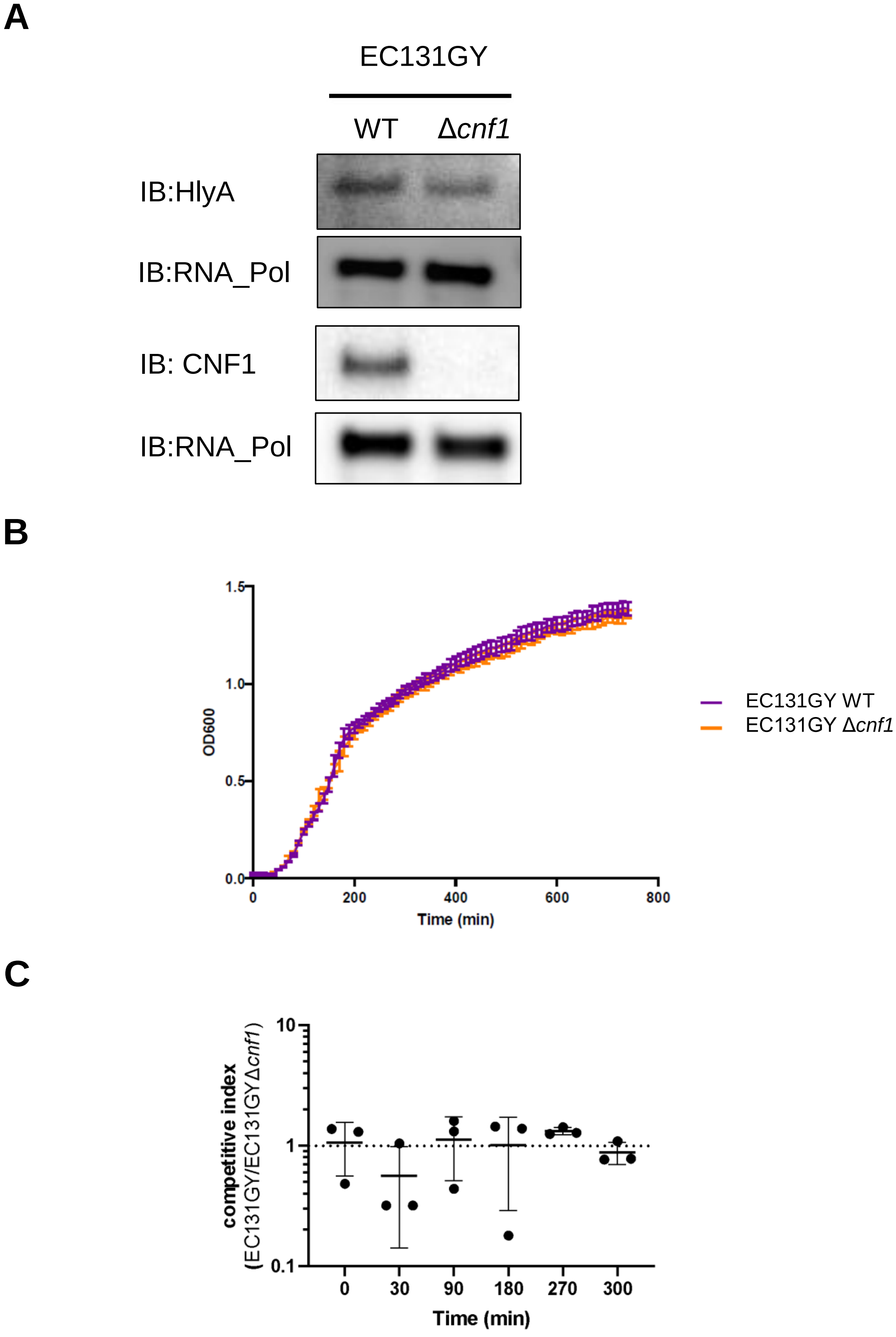


Sup. Fig. 5 : Tsoumtsa et al.
